# Aquatic metabolism throughout impoundment of a low productivity boreal reservoir using free water oxygen

**DOI:** 10.1101/2024.05.25.595904

**Authors:** D.P. Gedig, A.D. Soloway, R. Gill, T.N. Papakyriakou

## Abstract

Inland water bodies play a significant role in the global cycling of greenhouse gases (GHGs). Impoundment of rivers changes their GHG dynamics and leads to a pulse of emissions, primarily due to the respiration of introduced organic matter (OM). Aquatic metabolism estimates using free water oxygen curves were calculated at five sites on the lower Nelson River, Manitoba, Canada to assess the impact of impounding a new hydroelectric reservoir at the Keeyask Generating Station. After impoundment, the greatest ecosystem respiration (R) rates were seen in a former tributary inflow (-1.94 ± 1.03 g C m^-2^ d^-1^) and then in the forebay (-1.06 ± 0.74 g C m^-2^ d^-1^), an order of magnitude greater than the upstream extent of the reservoir area that appeared unimpacted (-0.13 ± 0.29 g C m^-2^ d^-1^). Benthic and pelagic R were of greater relative importance in the former tributary inflow and the forebay, respectively. Loading of allochthonous OM was a key factor regulating R. Further, evidence of “priming,” wherein labile OM facilitates the breakdown of more recalcitrant OM, was observed. Light limitation appeared to be prevalent throughout the study area, consistent with previous studies in the region. Despite the unique water chemistry present (i.e., high total phosphorus, high turbidity), aquatic metabolism in the study area appeared similar to other boreal impoundments. The results presented here updated the understanding of aquatic metabolism in a region characterized by hydroelectric development. Further, challenges in the methodology (e.g., gas transfer velocity estimation) were identified and discussed.

## Introduction

The importance of inland water bodies in the global carbon budget has recently been recognized. Previously thought of as a passive transport pathway from terrestrial to marine systems, surface inland waters play a significant role in both producing and metabolizing carbon (Cole et al. 2007; Regnier et al. 2022). Inland water bodies are mostly heterotrophic carbon emitters, and studying their dynamics is important toward better understanding drivers of global atmospheric greenhouse gas (GHG) concentrations (Cole et al. 2007; Duarte & Prairie 2005; Regnier et al. 2022). Examining anthropogenic influences on carbon cycling in aquatic systems, such as the impact of reservoirs, is an emerging field of study in the face of growing concerns surrounding climate change (Prairie et al. 2018; St. Louis et al. 2000; Tranvik et al. 2009). The combination of processes giving rise to carbon emissions in reservoir environments is complex, with pronounced spatiotemporal variability, depending on the characteristics of the reservoir, the contributing watershed, pre-flood landscape, productivity and climate (Deemer et al. 2016; Prairie et al. 2018; Rust et al. 2022). A pulse of increased GHG emissions (largely as CO_2_ and CH_4_) has often been documented after reservoir impoundment, attributed to the introduction of organic matter (OM) to the system after flooding and its subsequent respiration (Kelly et al. 1997; Rust et al. 2022; Teodoru et al. 2012). The net impact of reservoirs on GHG emissions requires an understanding of pre- and post-impoundment conditions and the impact of the reservoir on downstream biogeochemical water properties that give rise to GHG exchange (Maavara et al. 2020; Prairie et al. 2018). Investigating metabolism in a water body, specifically gross primary production (GPP) and ecosystem respiration (R), before and after impoundment provides direct insight into changes in carbon dynamics.

Both GPP and R influence the GHG balance in a water body through uptake and production of CO_2_, respectively (Duarte & Prairie 2005). Oxygen is consumed when CO_2_ is produced during respiration and vice versa during photosynthesis. As such, Odum (1956) was able to use information on changing dissolved oxygen concentrations ([O_2_]) with time to estimate metabolism in flowing water assuming that the rate of change in [O_2_] within a water volume were associated with three processes: GPP, R and air-water exchange (D), such that:

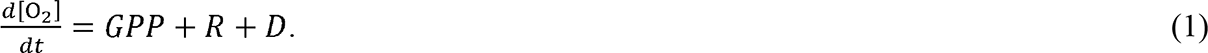

Some applications of Eq (1) involved bottle incubations (e.g., LaBuhn & Klump 2016), however, the use of “free water” oxygen curves provides estimates of metabolism within the total aquatic system rather than just the pelagic component (Pace & Prairie 2005; Pierson 2012). Due to advances in computing power and the ability to process lengthy time series datasets, the technique has produced metabolism estimates with increased accuracy (Appling et al. 2018a; 2018b; Winslow et al. 2016). Temporal and spatial comparison of calculated metabolism estimates would facilitate quantification of the changes in carbon dynamics caused by impoundment (e.g., respiration of introduced OM) and an understanding of how those changes vary throughout the reservoir area.

Increasingly, hydroelectric production of electricity is favourably looked on as a way to reduce GHG emissions in comparison to fossil fuels (Berga 2016). The proportion of electricity attributed to hydroelectric production varies by province within Canada. For example, in 2023 an estimated 97% of electricity generated in the province of Manitoba is in the form of hydroelectricity, compared to 63% in Canada as a whole (Canada Energy Regulator 2021).

Given this heavy reliance upon hydroelectric power in Manitoba, and more broadly in Canada, it is important to understand the impact of hydroelectric reservoirs on GHG emissions. The Keeyask Generating Station (KGS) is the most recent of a series of hydroelectric reservoirs constructed on the lower Nelson River in northern Manitoba (Fig. 1). Prior to construction, the Environmental Impact Statement estimated that one-half of the projected life-cycle GHG emissions of the KGS were attributable to land use changes, which were almost entirely due to reservoir impoundment (Keeyask Hydropower Limited Partnership [KHLP] 2012). The utility, Manitoba Hydro has monitored the physical and biogeochemical properties of the KGS site since 2009, covering the pre- and post-flood conditions. As part of this program, [O_2_] was continuously monitored during the ice-free season from within, as well as upstream and downstream of the KGS reservoir in 2020 and 2021, covering the first full year of impoundment. These data were used to calculate free water oxygen metabolism estimates, which were compared temporally and spatially to quantify the influence on biological CO_2_ dynamics in a boreal reservoir, with particular attention to increased respiration of introduced OM. Specifically, we assessed: (1) the immediate impact of impoundment on aquatic metabolism within the reservoir and downstream with consideration of variation upstream of the hydraulic zone of influence, (2) the range of GPP and R rates as observed in different sub-environments within the impounded reservoir and (3) the coupling of GPP and R using linear regression models. These results were then compared to previous studies on metabolism in natural waters and reservoirs to gain additional insight into the net impact of impoundment and how it may vary across reservoir space. In addition, challenges in the methodology for this particular environment were identified and improvements for possible future work were provided.

**Figure 1.**
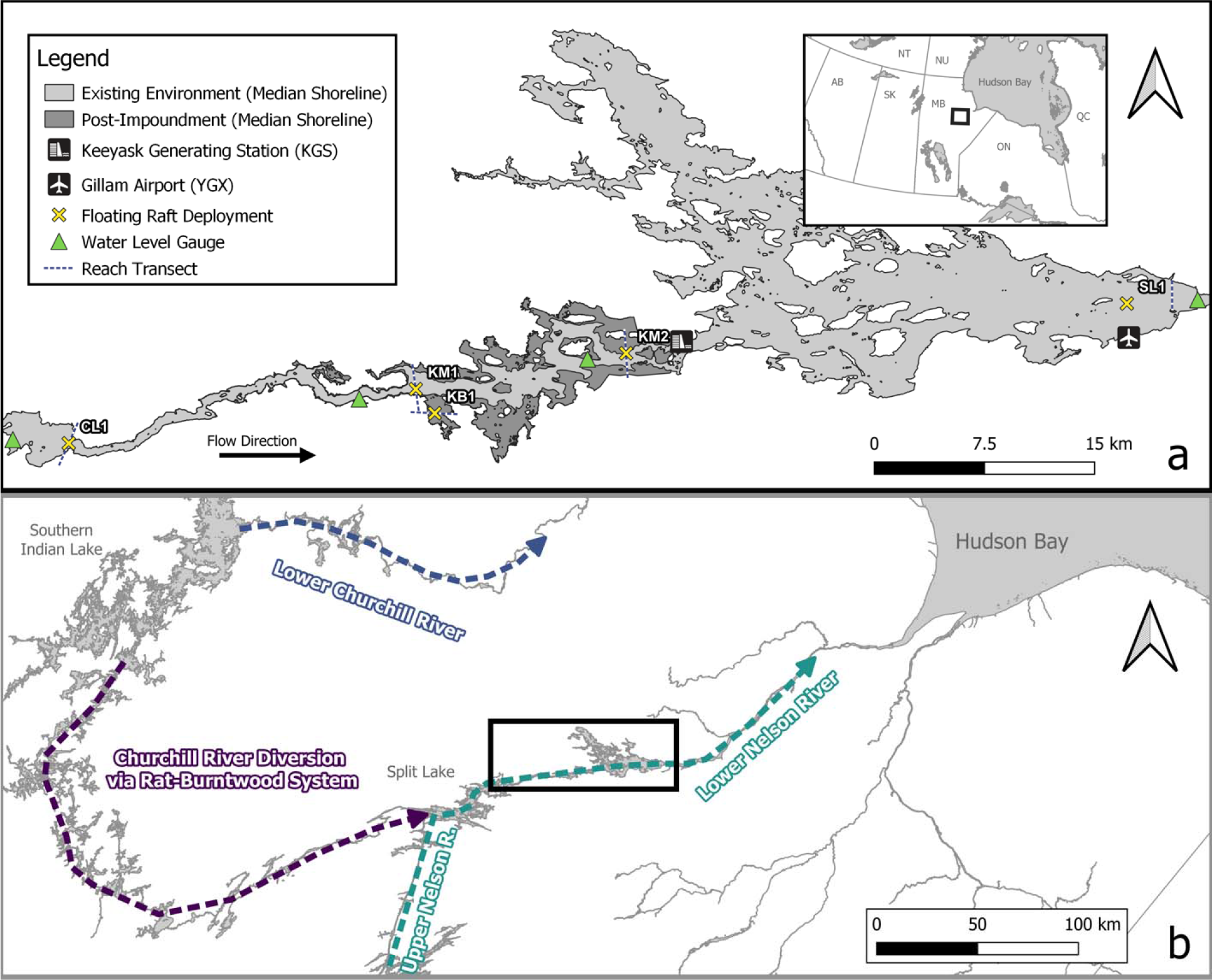
Map depicting the study area and operations occurring during 2020 and 2021. Floating raft IDs (three characters) are included. The black boxes in panel b and the inset map of panel a locate the study area. Created using QGIS (version 3.24.0).

## Methods

### Study area

The KGS is a 695-megawatt hydroelectric generating station located on the lower Nelson River in northern Manitoba, Canada (Fig. 1). Construction began in 2014, and the water-up process (refilling areas drained during construction) lasted from February 26 to April 20, 2020, with reservoir impoundment achieved between August 31 and September 5, 2020 (Manitoba Hydro 2021). The hydroelectric dams on the “lower” Nelson River (including KGS) receive flow from the Lake Winnipeg (the source) via the “upper” Nelson River as well as rerouted flow from an adjacent watershed associated with the Churchill River diversion (CRD) and downstream waters drain into Hudson Bay (Fig. 1b). Completed in 1976, the CRD impounded Southern Indian Lake and diverted flow from the Churchill River through the lakes comprising the Rat River and Burntwood River flowage (hereafter referred to as the Rat-Burntwood system) into the Nelson River (Fig. 1b; Newbury et al. 1984). The study area is in the Boreal Shield ecozone, with physiography consisting of peatlands overlying glaciolacustrine sediments above granitic and gneissic Precambrian bedrock of the Hudson Bay Lowlands (Betcher et al. 1995; Smith et al. 1998). The study area drainage is primarily organic soils (peatland forest interspersed with bog) with intermittent calcareous mineral soils, underlain by discontinuous permafrost (Smith et al. 1998). The median of daily mean discharge to the study area during the ice-free season (June to September) was 5444 m^3^ s^-1^ in 2020 and 2211 m^3^ s^-1^ in 2021, and only about 3% of the river flow is attributable to local inflow (KHLP 2012; Manitoba Hydro 2022). River water near the mainstem of the river is believed to be well mixed based on temperature data (KHLP 2012). The Nelson River has multiple tributary creeks within the study area that were expected to become wetland areas of longer hydraulic residence time (HRT) following impoundment, hereafter referred to as “backbays” (KHLP 2012). Based on limited continuous and discrete profile temperature and oxygen data from 2021, it appears that the backbay sampled in this study exhibits periodic stratification on the order of days (Manitoba Hydro 2022).

### Collection and sourcing of physicochemical data

Sensors deployed from floating rafts by the Manitoba Hydro Keeyask Physical Environment Monitoring Plan group (PEMP) at various locations throughout the study area continuously measured [O_2_] (mg L^-1^) and water temperature (°C) from late June (after ice break-up was complete) to September (sites described in Table 1; Fig. 1a) between 2020 and 2021. Monitoring operations were limited (particularly in 2020) due to complications related to the COVID-19 pandemic, resulting in some unavoidable incongruence between sampling sites and dates.

**Table 1.** The ID, decimal coordinates, location description and extent of metabolism data for each floating raft in the study area

| ID | Latitude<br>(deg) | Longitude<br>(deg) | Description | Data Extent<br>(DOY) |  |
| --- | --- | --- | --- | --- | --- |
|  |  |  |  | 2020 | 2021 |
| CL1 | 56.274 | -95.872 | Clark Lake; upstream of hydraulic zone of influence | 183 – 265 | 189 – 272 |
| KM1 | 56.315 | -95.493 | Entrance to Gull Lake; within (proposed) reservoir | 178 – 262 | 173 – 269 |
| KM2 | 56.342 | -95.263 | Keeyask Generating Station forebay; within reservoir | NA | 172 – 269 |
| KB1 | 56.301 | -95.471 | “Backbay” formed in tributary Rabbit Creek; within reservoir | NA | 171 – 269 |
| SL1 | 56.381 | -94.713 | Stephens Lake; downstream of reservoir | 179 – 231 | 174 – 268 |

Deployment dates were not exactly the same between sites or years but were considered comparable representations of the summer season (Table 1; Fig. S1). Measurement frequency was every 2 min in 2020 and every 5 min in 2021, except for KB1 which was measured every 10 min. Sensors were moored at a depth of 2 m below the surface, except for at site KB1 which was 0.5 m below the surface due to the shallower and less turbulent conditions. Discrete [O_2_] profiles taken on an approximately monthly basis during deployment were compared to validate measurements and confirm the continuous sensor is functioning properly. Continuous data are also screened annually with consideration of factors such as weather, hydroelectric infrastructure operation and water temperature, as well as compared to data from nearby sites. Due to a considerable discrepancy (> 1 mg L^-1^) in continuous [O_2_] measurements relative to discrete measurements throughout the season, [O_2_] data from SL1 in 2021 were reduced by 1 mg L^-1^ to address what appeared to be an error in calibration (further discussed in “Limitations, caveats and future work” subsection). The sensor assembly used for discrete profiles and most continuous measurements was the YSI EXO2 Multiparameter Sonde fitted with an EXO Optical Dissolved Oxygen Smart Sensor (Xylem Incorporated; ±0.1 mg/L or 1% of reading), but a HOBO Dissolved Oxygen Logger (Onset Computer Corporation; ±0.2 mg/L up to 8 mg/L, ±0.5 mg/L from 8 to 20 mg/L) was used for continuous measurements at site KB1. Calibrations were performed using air-saturated water as per manufacturer specifications.

From June to September in 2021, discrete water samples were collected on an approximately monthly basis by PEMP at 20% of the total water column depth using a water pump (Pentair Shurflo model 4008-101-E65) to fill two opaque 500 mL Nalgene bottles (filled to shoulder; for water chemistry parameters) and one 250 mL Pyrex bottle (filled with no headspace; for pH). All bottles were acid washed using 10% HCl (aq), rinsed with deionized water and triple-rinsed with sample water before use. Samples were kept chilled and in the dark during transport to the University of Manitoba (UM) and were usually received within 36 - 48 hours, but never more than 72 hours (dependent on delays related to the COVID-19 pandemic). Upon receipt at UM, water chemistry samples were filtered through pre-combusted Whatman Grade GF/C filters (1.2 µm nominal pore size) for analysis at the certified water chemistry laboratory at the Freshwater Institute, Department of Fisheries and Oceans (Winnipeg, MB) as per Stainton et al. (1977).

Samples were analyzed for dissolved organic carbon (μmol L^-1^), chlorophyll a (μg L^-1^), total nitrogen (reported as sum of total dissolved nitrogen and suspended nitrogen; μg L^-1^) and total phosphorus (reported as sum of total dissolved phosphorus and suspended phosphorus; μg L^-1^). Filtrate (kept at 4°C) and filters were submitted once the latter had been desiccated for at least 24 hours. The pH of each sample was measured at UM in lab conditions (∼20.5°C) using an Orion Star 121 (± 0.01 pH units) after a three-point calibration with temperature adjustment (pH 4.01, 7.00 and 10.01 pH at 25°C buffer solutions).

Photosynthetic photon flux density (PPFD; µmol m^-2^ d^-1^) data were obtained from the NASA Langley Research Center (LaRC) POWER Project funded through the NASA Earth Science/Applied Science Program. Hourly data of barometric pressure and wind speed at 10 m from the Gillam Airport (YGX) were extracted from the Environment and Climate Change Canada Historical Climate Data website (https://climate.weather.gc.ca/index_e.html) throughout 2022. Missing wind speed and pressure data from YGX was replaced with the daily mean. Mean depth (z_J) was calculated using water level data (m a.s.l.) from gauges throughout the study area and bathymetry data from Manitoba Hydro (Fig. 1a). Water level data were visually screened for errors before use and outlier values were removed. Water level data was linearly interpolated to the temporal resolution of [O_2_] measurements before use. Flow path of the Nelson River mainstem was determined by generating a Voronoi diagram and tracing the centerline bisecting the river with minor manual smoothing (i.e., vertex removal) considering prior knowledge of flow (KHLP 2012). The transect of the reach was established as a line perpendicular to the centerline and that intersected the nominal coordinates of each floating raft. The centerline and all transects were determined using post-impoundment conditions for consistency. The elevation profile of the reach at each floating raft was determined using the 3D Analyst extension in ArcGIS Pro (version 3.0.3) and linearly interpolated. The z_J was calculated as the cross-sectional area divided by the reach length at a given water level. Bathymetric data used in interpolations and other calculations maintained a minimum resolution 0.001 m. Due to limited data in Stephens Lake, depth measurements from a transect 3 km downstream of the floating raft were used to estimate the actual reach profile for SL1. The resultant elevation profile was multiplicatively scaled in both dimensions to reflect actual site depth and reach length before usage in calculations.

### Model structure and data preparation

The open-source R package *LakeMetabolizer* was used to calculate daily estimates of metabolism, in which GPP is positive and R is negative by convention (Winslow et al. 2016). Measurements of [O_2_] were smoothed using a rolling mean over 100 min to reduce noise.

Missing or omitted data were replaced with the last available observation, and days with more than 20 min of missing or omitted data were removed from the time series. The equilibrium saturated [O_2_] was calculated after Garcia and Gordon (1992) assuming salinity of zero. Gas transfer velocity (k in m d^-1^) normalized to a Schmidt number of 600 (k_600_) was estimated hourly using wind speed and surface area (km^2^) after Vachon and Prairie (2013) and scaled based on water temperature to the temporal resolution of the [O_2_] measurements (Jähne et al. 1987; Raymond et al. 2012). We opted for k_600_ estimation based on wind-driven turbulence as it becomes a primary process in larger rivers (Hall & Ulseth 2020) and is used in similar applications (e.g., Demarty et al. 2009). The *steamMetabolizer* R package was used to interpolate PPFD observations through merging with modelled curves based on geographic location (Appling et al. 2018a). We assumed the water column was entirely mixed, which may have been partially violated at KB1 because of periodic stratification but was considered tolerable given the handling of process errors (discussed below). The volumetric estimates of NEP produced by the model (mg O_2_ L^-1^ d^-1^) were depth integrated by multiplying by the daily mean of z_J at each site and converted into equivalent quantities of carbon (g C m^-2^ d^-1^) assuming a photosynthetic quotient (PQ) of 1.2 and respiratory quotient (RQ) equal to the inverse, which closely reflects Yezhova et al. (2021). Areal oxygen-based NEP estimates obtained from other studies were converted using the PQ and/or RQ values chosen by the authors if provided.

Free water oxygen metabolism calculation may produce theoretically impossible estimates when unconstrained (i.e., negative GPP or positive R), attributed to physical processes occluding the biological signal (Rose et al. 2014; Winslow et al. 2016). An approach to handling outlier values is to retain them and interpret the mean estimate (Staehr et al. 2010), as is adopted here. The *LakeMetabolizer* model option initially used was maximum likelihood estimation (MLE) including process error only (Solomon et al. 2013; Winslow et al. 2016). Despite the strength afforded by the longer time series here, CL1 exhibited mean impossible values when using MLE which invalidates any interpretation of GPP or R. For this site only, the “bookkeeping” model option was used instead for both years, which includes no error terms and does not fit estimates using environmental variables. This option still produces a valid NEP estimate when assuming all variation in the [O_2_] signal outside of D is attributable to biological processes. Further, NEP estimates are generally considered more robust than gross metabolic rates (Chapin et al. 2006; Staehr et al. 2010). Photochemical oxidation has been found to be a notable influence in high latitude freshwater carbon dynamics (Cory et al. 2014), which may have introduced error into metabolism estimates but is not separately discussed here.

### Statistical analyses

Pairwise two-tailed Welch’s t-tests were performed to test for differences in volumetric estimates of NEP using data from sites measured in both 2020 and 2021. Due to data loss at SL1 in 2020, the temporal t-test for this site was performed using only data before day-of-year 232 for both years. A one-way Welch’s Analysis of Variance (ANOVA) was performed to test for differences in volumetric and areal estimates of NEP from 2021. Differences in GPP and R estimates from within the impounded reservoir in 2021 were also tested using a one-way Welch’s ANOVA. Post hoc analysis was performed using a Games-Howell multiple comparisons test and results were reported using compact letter display (CLD; Piepho 2004). Although consequences from violating the assumption of normality is minimized if sample sizes are > 40 (as is true in all cases in this study), normality was examined by visual assessment of density plots (Ghasemi & Zahediasl 2012). The critical *P*-value for statistical tests was α = 0.05. Coupling of R and GPP was examined using regression models as in Solomon et al. (2013) to determine the level of “background respiration” (β_0_; respiration of allochthonous and recalcitrant autochthonous OM) and the slope of the relationship (β_1_; equal to one when ratio of units produced to units respired is 1:1). Briefly, models were fit including a first-order autoregressive term after standardizing R and GPP values to 20°C (Holtgrieve et al. 2010), and the β_0_ and β_1_ coefficients were qualitatively interpreted in the context of the metabolism results (Eq (2))

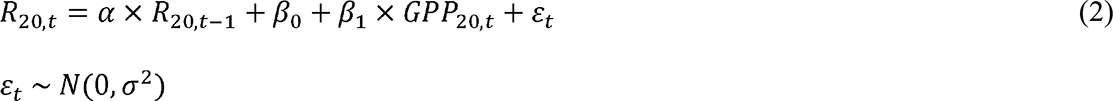

A first-order autoregressive model was implemented for simplicity and consistency with Solomon et al. (2013). To estimate uncertainty in β_0_ and β_1_, 95% confidence intervals were calculated. Statistical analyses were performed using R (version 4.2.2) within RStudio (version 2023.3.0.386) and graphics were created using the *ggplot2* package (R Core Team 2022; RStudio Team 2020; Wickham 2016).

## Results

In 2020, the lowest median [minimum - maximum] percent [O_2_] saturation was observed at KM1 (101.6% [75.4 - 107.6%]), followed by CL1 (104.9% [101.2 - 110.3%]) and then SL1 (107.6% [101.6 - 119.2%]). The greatest median water temperature was observed at SL1 (19.71° [14.03 - 22.21°]), followed by CL1 (18.76° [9.47 - 22.07°]) and then KM1 (18.73° [8.17 - 22.15°]). In 2021, the median percent [O_2_] saturation in ascending order was KB1, KM2, KM1, CL1 then SL1 (Table 2). Median water temperature in descending order was CL1, KM1, SL1, KM2 then KB1 (Table 2). Across the study area, the median dissolved organic carbon (DOC) concentration was 745.5 µmol L^-1^ [658 – 902 µmol L^-1^] (Fig. 2a). The median chlorophyll a concentration was 1.46 µg L^-1^ [0.77 - 24.26 µg L^-1^] (Fig. 2b). The median total dissolved and suspended nitrogen concentrations were 411.5 µg L^-1^ [358 - 463 µg L^-1^] and 40.5 µg L^-1^ [16 - 255 µg L^-1^], respectively (Fig. 2c). The median total dissolved and suspended phosphorus concentrations were 36.5 µg L^-1^ [24 - 55 µg L^-1^] and 8.5 µg L^-1^ [5 - 35 µg L^-1^], respectively (Fig. 2d). The waters studied here were alkaline with a median lab pH of 7.87 [7.45 - 8.01]. When classified using the Trophic State Index of Carlson and Simpson (1996), the study area was generally oligomesotrophic based on chlorophyll a concentration but exhibited total phosphorus concentrations associated with eutrophy (Fig. 2).

**Figure 2.**
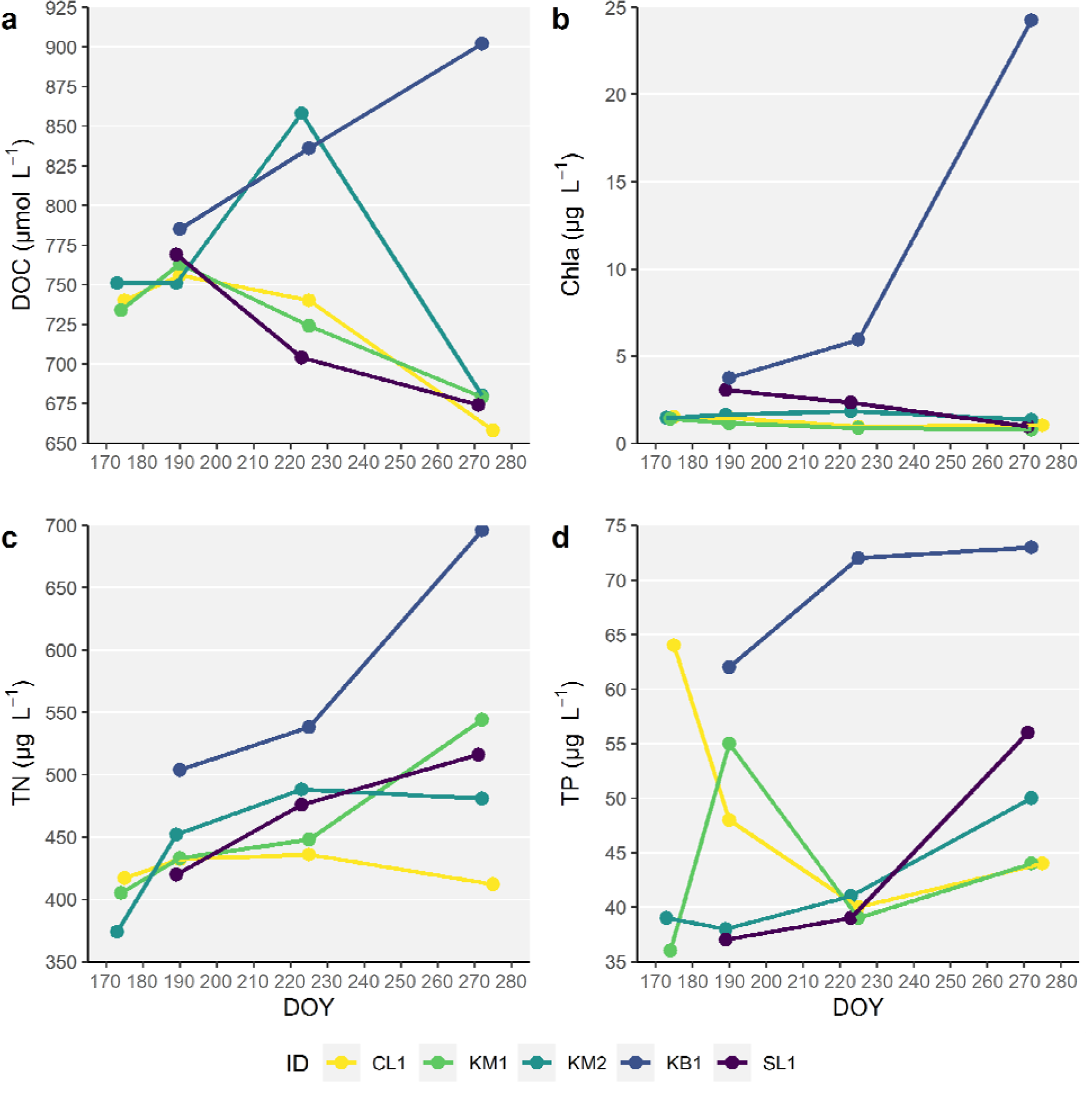
Time series plots of dissolved organic carbon (a), chlorophyll a (b), total nitrogen (c) and total phosphorus (d) at sites near and within the Keeyask Generating Station reservoir in northern Manitoba, Canada during 2021. Samples were collected by the Manitoba Hydro Physical Environment Monitoring Plan group and analyzed at the Freshwater Institute, Department of Fisheries and Oceans.

**Table 2.** Univariate summary of physicochemical parameters as median [minimum - maximum] and aquatic metabolism estimates as mean (± standard deviation) for each site in the study area during 2021.

|  | CL1 | KM1 | KM2 | KB1 | SL1 |
| --- | --- | --- | --- | --- | --- |
| **z**<br>(m) | 2.34<br>[2.29 - 2.78] | 10.39<br>[10.18 - 10.52] | 7.18<br>[7.02 - 7.26] | 2.34<br>[2.27 - 2.36] | 18.16<br>[17.24 - 18.67] |
| **k**<br>(m d<sup>-1</sup>) | 2.13<br>[0.69 - 6.99] | 2.43<br>[0.68 - 8.19] | 2.47<br>[0.55 - 8.12] | 2.49<br>[0.55 - 7.8] | 2.64<br>[0.55 - 8.91] |
| **T**<br>(°C) | 17.11<br>[11.64 - 20.84] | 17.00<br>[11.92 - 20.81] | 16.64<br>[11.61 - 20.81] | 16.42<br>[10.1 - 21.78] | 16.89<br>[11.99 - 20.16] |
| **[O<sub>2</sub>]**<br>(% sat.) | 98.7<br>[94.3 - 103.7] | 98.1<br>[94.4 - 103.1] | 94.2<br>[82.6 - 104.3] | 84.8<br>[56.2 - 106.6] | 100.1<br>[95.8 - 114.9] |
| **R**<br>(mg O<sub>2</sub> L<sup>-1</sup> d<sup>-1</sup>)<br>(g C m<sup>-2</sup> d<sup>-1</sup>) | NA<br>NA | -0.041 (± 0.088)<br>-0.13 (± 0.29) | -0.47 (± 0.33)<br>-1.06 (± 0.74) | -2.66 (± 1.40)<br>-1.94 (± 1.03) | NA<br>NA |
| **GPP**<br>(mg O<sub>2</sub> L<sup>-1</sup> d<sup>-1</sup>)<br>(g C m<sup>-2</sup> d<sup>-1</sup>) | NA<br>NA | 0.0042 (± 0.0531)<br>0.014 (± 0.173) | 0.28 (± 0.25)<br>0.63 (± 0.56) | 0.72 (± 0.79)<br>0.52 (± 0.58) | NA<br>NA |
| **NEP**<br>(mg O<sub>2</sub> L<sup>-1</sup> d<sup>-1</sup>)<br>(g C m<sup>-2</sup> d<sup>-1</sup>) | -0.12 (± 0.29)<br>-0.089 (± 0.207) | -0.037 (± 0.067)<br>-0.12 (± 0.22) | -0.19 (± 0.13)<br>-0.43 (± 0.30) | -1.95 (± 1.32)<br>-1.42 (± 0.97) | 0.012 (± 0.128)<br>0.069 (± 0.724) |

Mean volumetric NEP was significantly different between 2020 and 2021 at CL1 (*P* < 0.001), KM1 (*P* = 0.001) and SL1 (*P* = 0.002; Fig. 3). The difference (mean ± SE) was greatest at CL1 (-0.45 ± 0.04), followed by KM1 (-0.07 ± 0.02) and SL1 (-0.07 ± 0.02). There were significant differences in mean volumetric and areal NEP (*P* < 0.001 both) between sites in 2021 (Table 2; Fig. 4; Fig. 5 with CLDs presented). Site KB1 had the lowest NEP on both a volumetric and areal basis, followed by KM2 (Table 2; Fig. 4; Fig. 5). Between sites within the KGS reservoir, there were significant differences in GPP and R on a volumetric and areal basis (*P* < 0.001 all; Table 2; Fig. 6; Fig. 7 with CLDs presented). Site KB1 had the greatest volumetric and areal R, followed by KM2 and then KM1 (Table 2; Fig. 6; Fig. 7). The greatest volumetric GPP was at KB1, followed by KM2 and then KM1 (Table 2; Fig. 6). Areal GPP was not significantly different between KB1 and KM2, but was lowest at KM1 (Table 2; Fig. 7).

**Figure 3.**
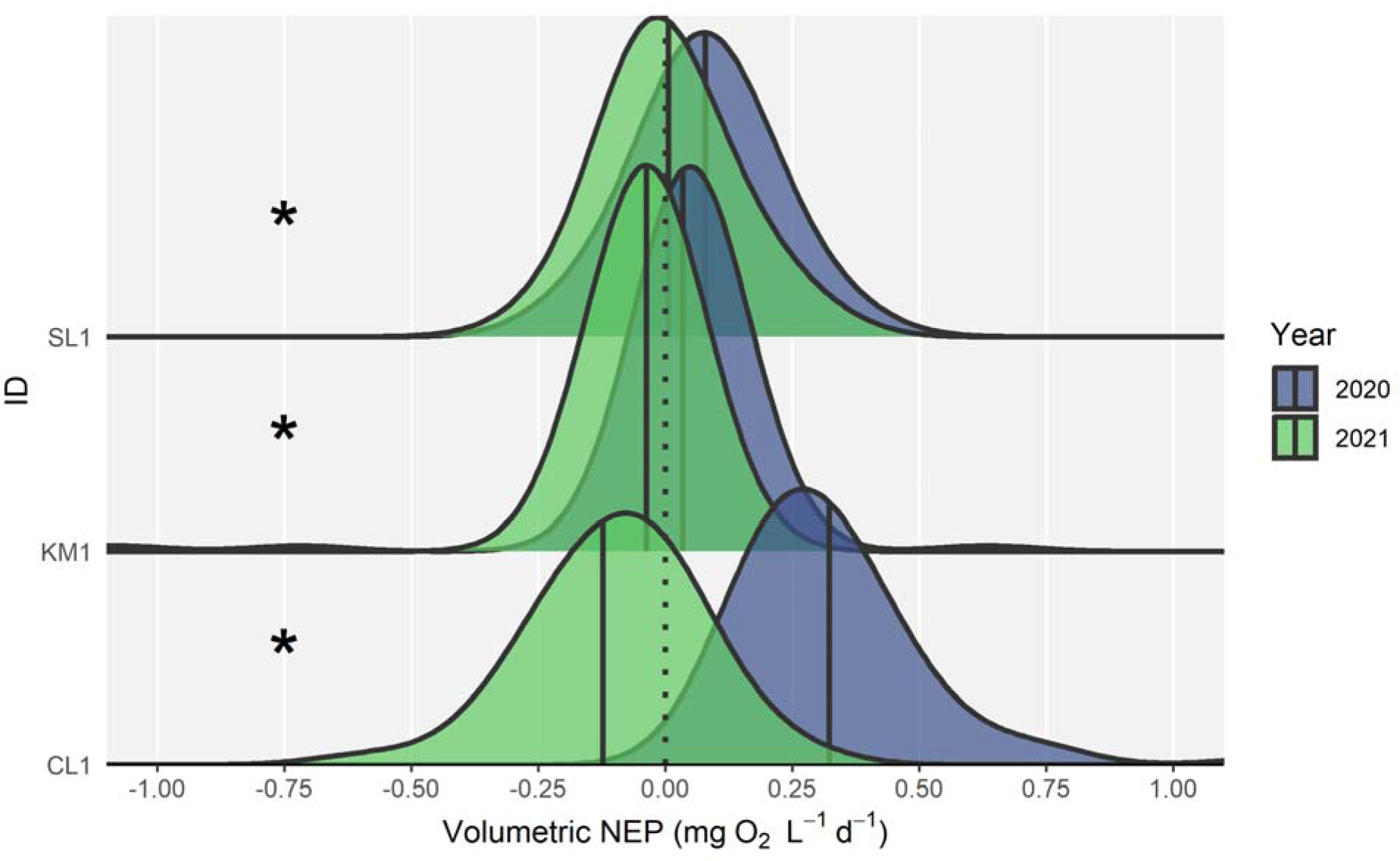
Density plots of volumetric net ecosystem production (NEP in mg O_2_ L^-1^ d^-1^) by site near and within the Keeyask Generating Station reservoir in northern Manitoba, Canada. An asterisk indicates significant difference between years via Welch’s two-sample t-test (α = 0.05). The mean of each distribution is represented by a solid vertical line. Data presented in this figure and used in the t-test for SL1 is from days before day-of-year 232.

**Figure 4.**
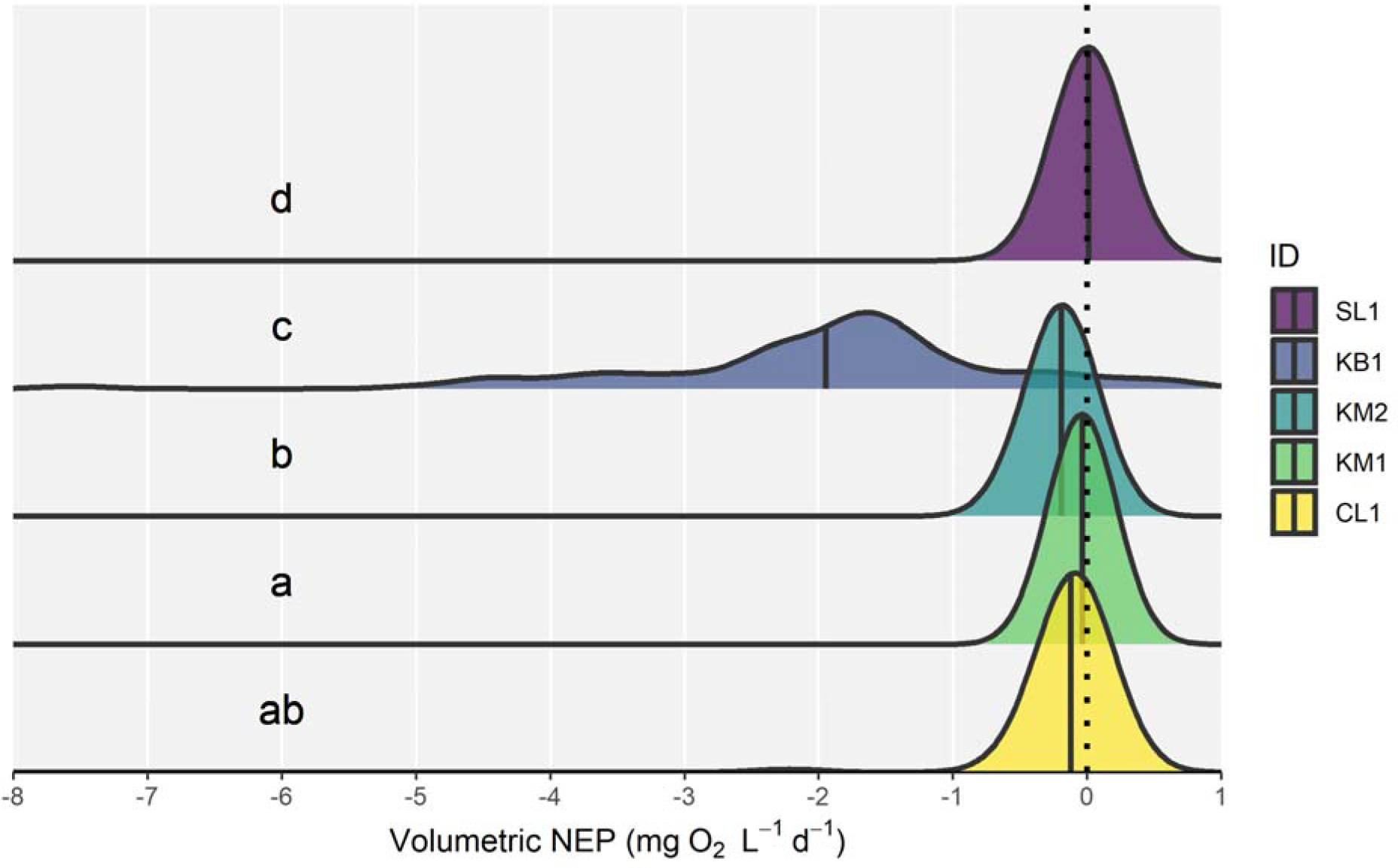
Density plots of volumetric net ecosystem production (NEP in mg O_2_ L^-1^ d^-1^) by site near and within the Keeyask Generating Station reservoir in northern Manitoba, Canada during 2021. The mean of each distribution is represented by a solid vertical line. Results of Games-Howell post-hoc test are presented using compact letter display.

**Figure 5.**
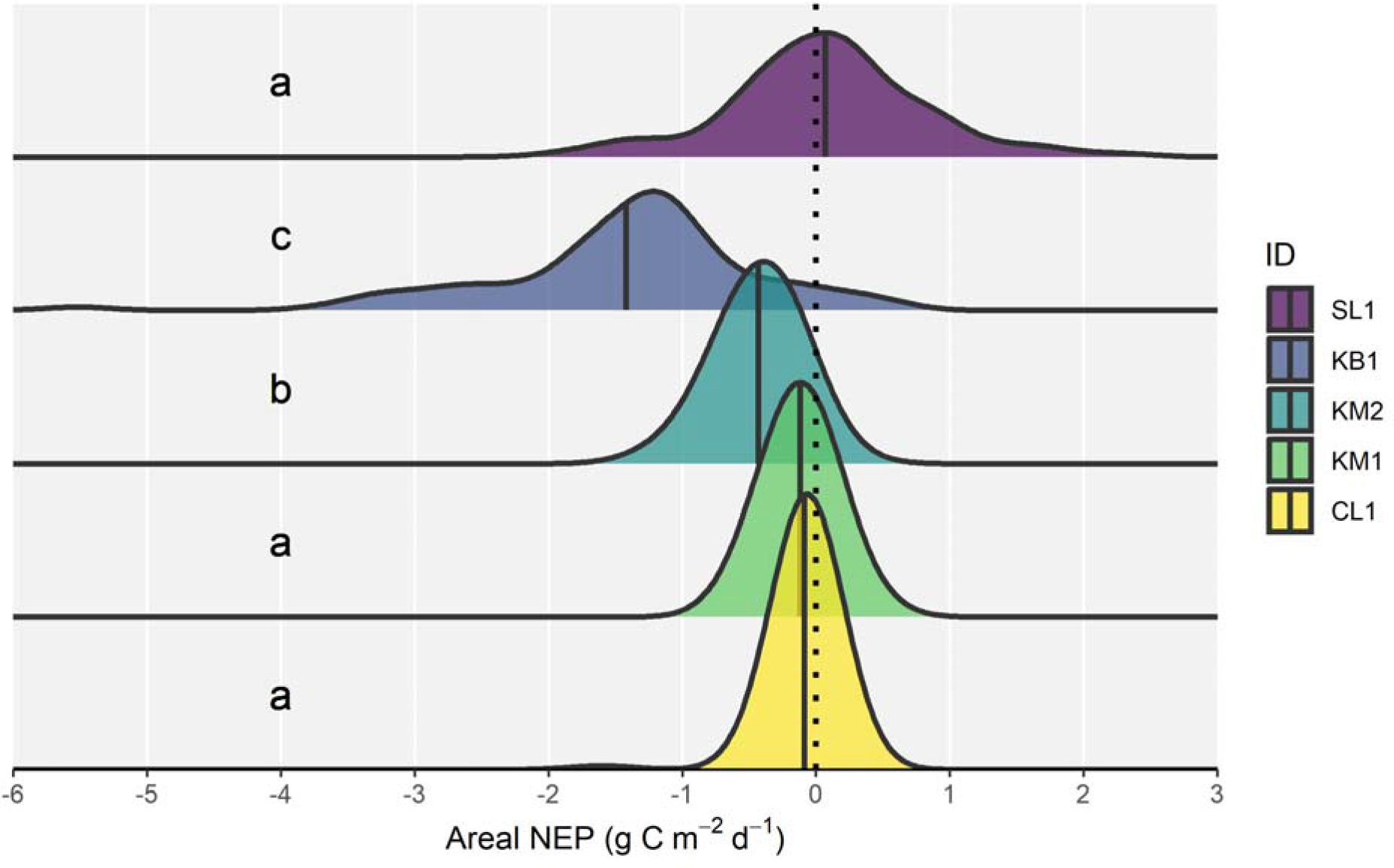
Density plots of areal net ecosystem production (NEP in g C m^-2^ d^-1^) by site near and within the Keeyask Generating Station reservoir in northern Manitoba, Canada during 2021. The mean of each distribution is represented by a vertical solid line. Results of Games-Howell post-hoc test are presented using compact letter display.

**Figure 6.**
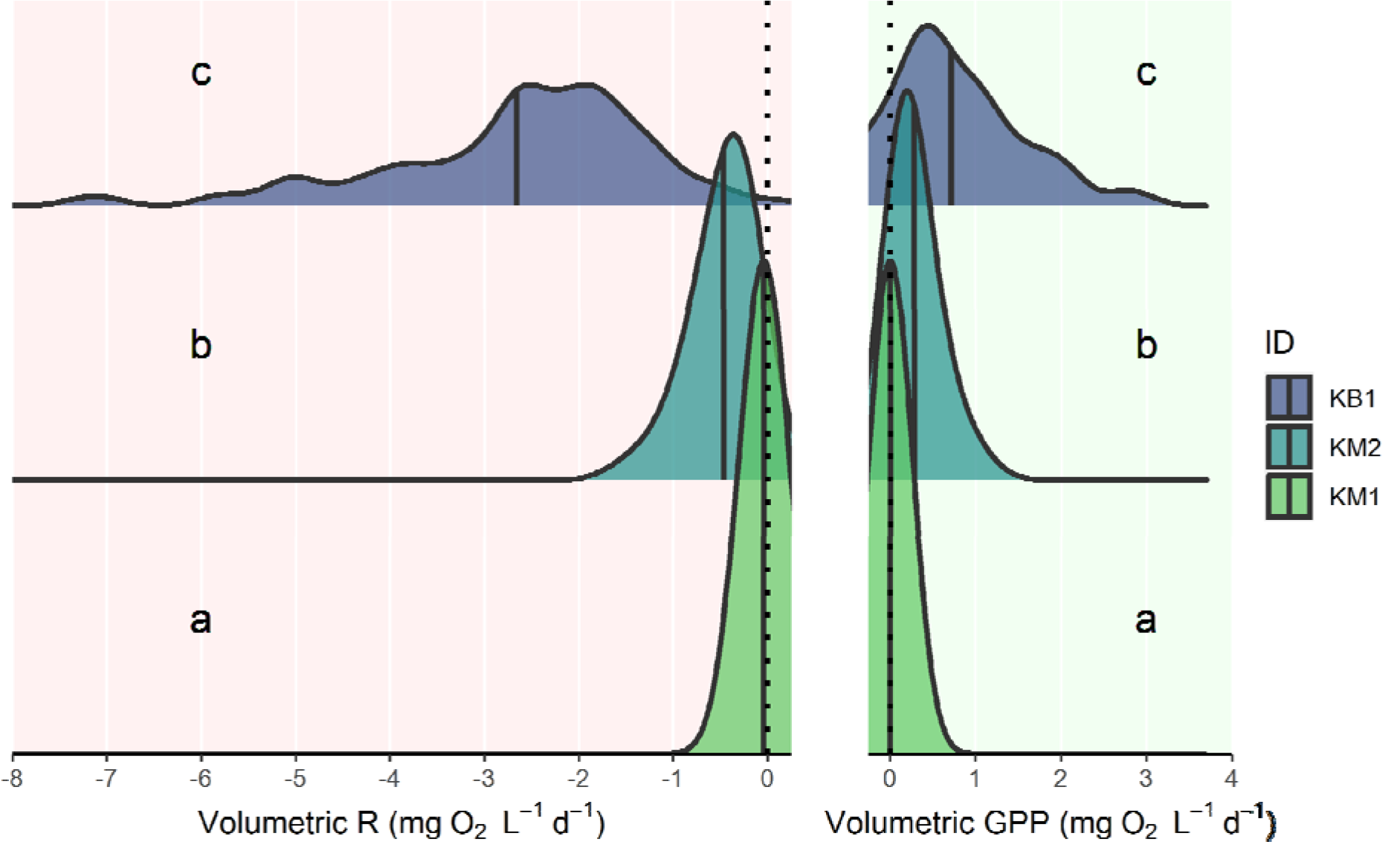
Density plots of volumetric ecosystem respiration (R in mg O_2_ L^-1^ d^-1^) and gross primary production (GPP in mg O_2_ L^-1^ d^-1^) by site within the Keeyask Generating Station reservoir in northern Manitoba, Canada during 2021. The mean of each distribution is represented by a vertical solid line. Results of Games-Howell post-hoc test are presented using compact letter display.

**Figure 7.**
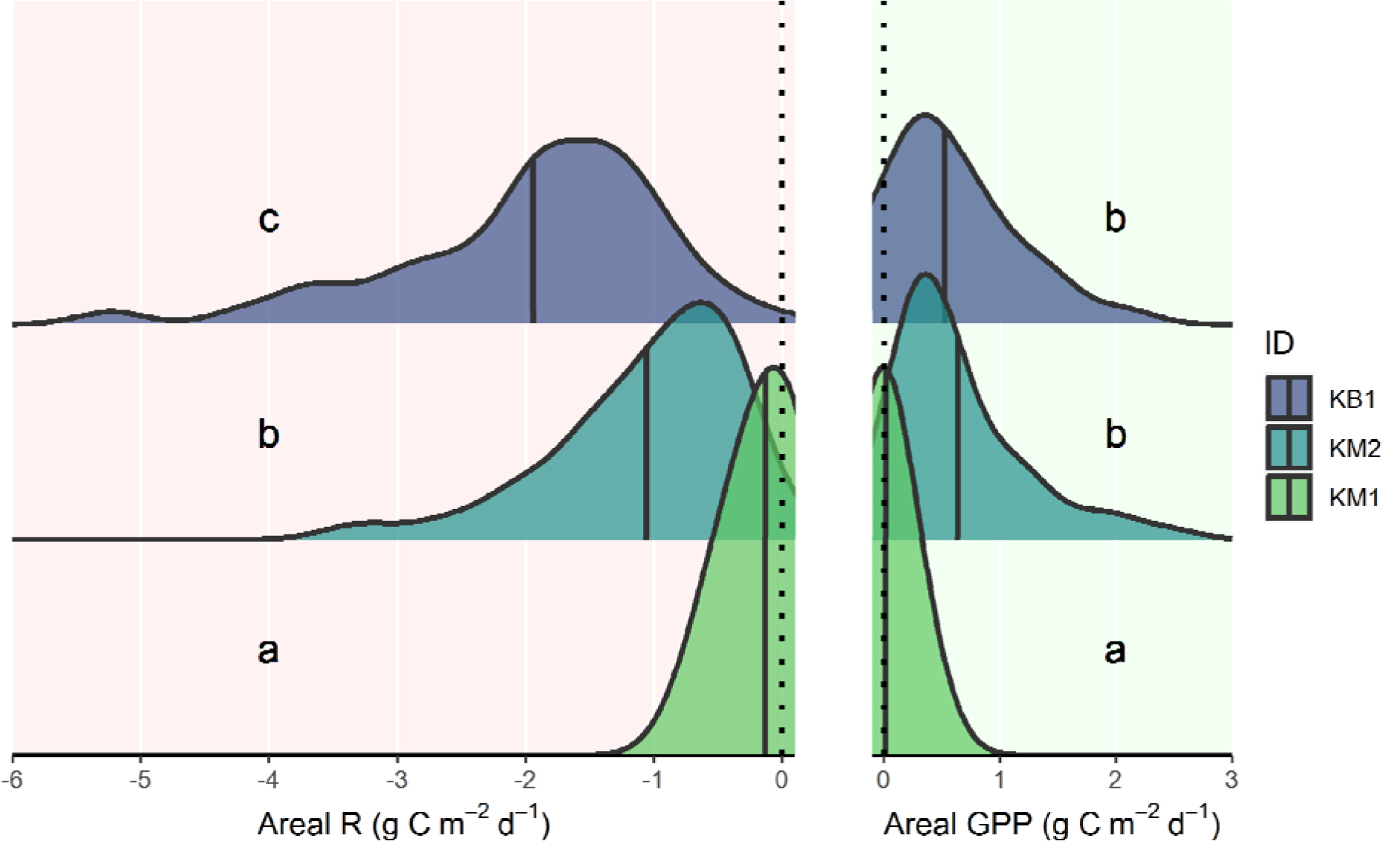
Density plots of areal ecosystem respiration (R in g C m^-2^ d^-1^) and gross primary production (GPP in g C m^-2^ d^-1^) by site within the Keeyask Generating Station reservoir in northern Manitoba, Canada during 2021. The mean of each distribution is represented by a vertical solid line. Results of Games-Howell post-hoc test are presented using compact letter display.

Based on autoregressive linear modeling, KB1 had the lowest β_0_ value or highest level of background respiration, followed by KM2 and then KM1 (Table 3; Fig. 8). Site KM2 had the greatest β_1_ value or exhibited the greatest degree of R and GPP coupling, followed by KM1 and then KB1 (Table 3; Fig. 8). The 95% confidence margins of error for β_0_ and β_1_ did not overlap 0 or 1 for any site (Table 3).

**Figure 8.**
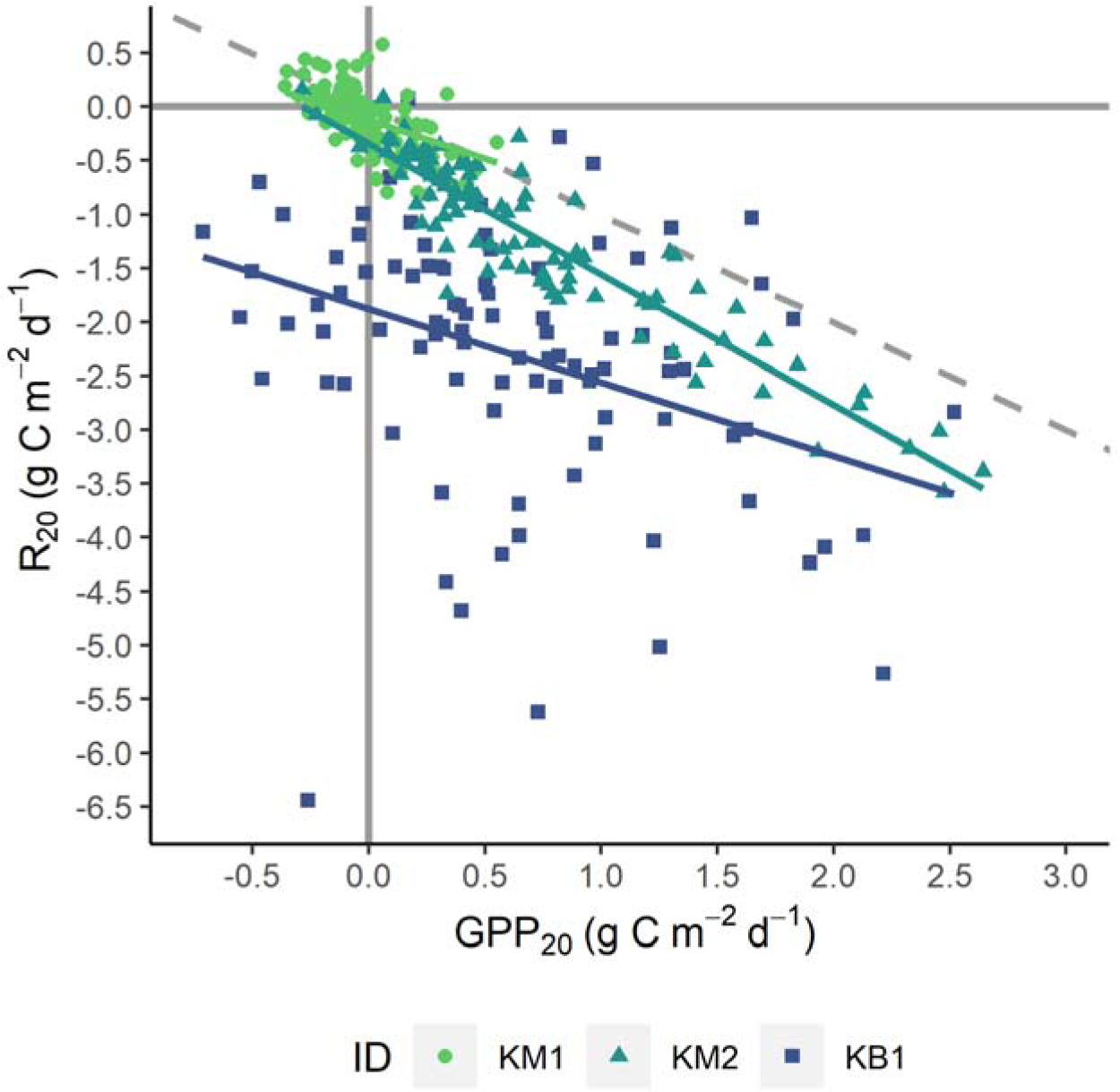
Ecosystem respiration versus gross primary production adjusted to 20°C (R_20_ and GPP_20_ respectively in g C m^-2^ d^-1^) by site within the Keeyask Generating Station reservoir in northern Manitoba, Canada during 2021. A -1:1 line from the origin (dashed) and first-order autoregressive linear relationships (solid) are shown.

**Table 3.** Fitted coefficients with 95% confidence margins of error and *P*-values from autoregressive linear modeling of ecosystem respiration versus gross primary production adjusted to 20°C (g C m^-2^ d^-1^). No *P*-value is provided for the α coefficient.

| ID | $\alpha$ | $\beta_0$ (g C m<sup>-2</sup> d<sup>-1</sup>) | | | *P* | $\beta_1$ | | | *P* |
| --- | --- | --- | --- | --- | --- | --- | --- | --- | --- |
| KB1 | 0.39 | -1.88 | ± | 0.04 | < 0.001 | -0.68 | ± | 0.03 | < 0.001 |
| KM1 | 0.61 | -0.12 | ± | 0.01 | < 0.001 | -0.72 | ± | 0.02 | < 0.001 |
| KM2 | -0.16 | -0.35 | ± | 0.01 | < 0.001 | -1.21 | ± | 0.01 | < 0.001 |

## Discussion

### Spatial and temporal variability in aquatic metabolism

Locations above, within and downstream of the KGS reservoir were all significantly more heterotrophic immediately following impoundment, and the comparable effect observed upstream confounds the interpretation of its metabolic impact. The NEP estimates at mainstem sites (as those used in the temporal comparison) appeared subject to greater error than the backbay when compared to estimates calculated using dissolved CO_2_ (Fig. S3). These errors were likely due to physical processes dominating the [O_2_] signal (Rose et al. 2014), and turbulence at these locations may have yielded k values poorly described by published models (Hall & Ulseth 2020). As a consequence, these temporal differences were difficult to interpret despite being statistically significant, and the deviations from zero mean NEP were likely negligible. Site KM1 was at the upstream extent of the reservoir and may have received less of the introduced nutrient load, in contrast to the downstream KM2 in the forebay that was significantly more heterotrophic. For reservoirs that maintain riverine characteristics such as the KSG, it seems selecting locations where the effects of impoundment will be most pronounced is key for their accurate assessment. If one anticipates that k will be difficult to constrain due to strong physical processes, usage of alternative empirical approaches (e.g., calculating k_600_ using eddy covariance and environmental parameters) is advised, especially in areas with low productivity (Hall & Ulseth 2020).

Overall, the impounded reservoir was heterotrophic on average, which is consistent with prior study of inland waters (Bogard et al. 2020; Cole et al. 2007). The backbay and forebay were the most heterotrophic sub-environments, and likely exhibited the greatest CO_2_ emissions given their association with respiration in boreal waters (Barbosa et al. 2023; Vachon et al. 2017), especially considering how internally produced CO_2_ comprises a greater proportion of emissions in larger rivers (Hotchkiss et al. 2015). The backbay sustained higher DOC concentration into the late summer in contrast to other sites, which is associated with greater levels of respiration and primary production (Bogard et al. 2020; Hanson et al. 2003). This is consistent with how reservoir construction increases nutrient retention and mineralization of organic carbon, increasingly so with longer HRT (Maavara et al. 2017; 2020). Benthic respiration is a larger share of the total aquatic respiration in shallower littoral areas, which may also influence why the relatively shallow backbay is overwhelmingly the most heterotrophic on a volumetric basis (Pace & Prairie 2005). After impoundment of the reservoir directly downstream the KGS (Stephens Lake), primary productivity was lower in a backbay than the mainstem despite elevated nutrient concentrations (Hecky & Harper 1974). This inhibition was attributed to nutrient limitation after iron was complexed with introduced OM and rendered biologically unavailable (Guildford et al. 1987; Jackson & Hecky 1980). In the KGS reservoir, the greater volumetric primary production observed in the backbay versus mainstem locations does not reflect such inhibition.

Within the KGS reservoir, autoregressive linear modeling indicated that respiration and primary production coupling was greater than 1:1 in the forebay, whereas coupling was less than 1:1 in the backbay and the upstream mainstem. The tight coupling of respiration and primary production in oligotrophic lakes appears to be from substrate limitation of heterotrophs, whereas the uncoupling observed in more eutrophic lakes may be due to production rates exceeding respiration capacity (Sadro et al. 2011; Solomon et al. 2013). This suggests there is a quasi- eutrophic state in the backbay wherein some degree of autochthonously produced OM escapes mineralization, perhaps in part due to supplementation of allochthonous OM as evidenced by the comparatively high level of background respiration. Uncoupling in the upstream mainstem location is harder to explain given the low rates of gross metabolic processes, and may be due to factors such as nutrient level does not seem to control metabolism in flowing waters as it does in lakes (Bernhardt et al. 2018). The greater than 1:1 coupling observed in the forebay could be a result of “priming”, in which the availability of labile autochthonous OM facilitates the degradation of more recalcitrant OM (Guenet et al. 2010; Solomon et al. 2013). Sources of particulate OM in previously impounded areas of the lower Nelson River are predominantly proximal (i.e., from locally-derived soils/sediments and tributaries; Stainton 2019), and priming could promote mineralization (rather than sedimentation or export) of introduced terrestrial OM that is generally considered recalcitrant (Guenet et al. 2010). As observed in permafrost soils exhibiting characteristic cryoturbation, the OM mobilized from near surface permafrost may be more biodegradable than that of the active layer (Diochon et al. 2013). More broadly, permafrost soils exhibited a greater proportion of readily biodegradable OM than soils without permafrost in a meta-analysis by Vonk et al. (2015). Previous study suggests the quantity of labile OM exported to larger rivers decreases across the summertime (Palviainen et al. 2022; Vonk et al. 2015). This aligns with the declining DOC concentrations at seemingly unimpacted locations seen here, and is supported by the high biodegradability of lower Nelson River OM in wintertime documented by Kazmiruk et al. (2021). In contrast, both the forebay and the backbay exhibited relatively elevated DOC concentration later into the summer, suggesting continued loading of OM.

### Comparison to other relevant ecosystems

The Nelson River has historically had lower primary production in comparison to the Rat-Burntwood and Churchill River systems, attributed to higher turbidity causing light limitation (Hecky & Harper 1974). After impoundment of Southern Indian Lake as part of the CRD, mainstem areas of the Rat-Burntwood system exhibited increased turbidity in response to eroding fine-grained glacio-lacustrine deposits, leading to a light limitation that relieved phosphorus deficiency but did not alter primary production on an areal basis (Guildford et al. 1987; Hecky & Guildford 1984). The lack of change in areal primary production was attributed to compensatory adaptation of the phytoplankton community (e.g., higher chlorophyll concentrations, increased photosynthetic efficiency per unit of chlorophyll; Hecky & Guildford 1984). Areas outside of the mainstem flow of the Rat-Burntwood system (comparable to backbays) that did not experience increases in suspended sediment appeared to instead be stimulated by the introduced nutrients (Hecky & Guildford 1984). Here, the high phosphorus concentrations in the study area are reminiscent of this light limitation, and the late-season spike of chlorophyll a concentration in the backbay alongside the non-significant difference in primary production between the detectably impacted sub-environments suggests analogous accommodation to varying conditions. Erosional processes (e.g., bank degradation, permafrost thaw) in the Rat-Burntwood system and locally caused increases in total suspended sediment concentration within prior impoundments on the lower Nelson River (Stainton 2019).

Lake Winnipeg exhibited a mean GPP of 1.1 ± 0.9 g C m^-2^ d^-1^ and mean R of -8.1 ± 6.0 g C m^-2^ d^-1^ in 2018, and was net heterotrophic overall (Yezhova et al. 2021). Here, primary production was only comparable to Lake Winnipeg in the forebay and backbay (i.e., impacted sub-environments), and respiration rates appear lower than Lake Winnipeg on average even in the most heterotrophic areas. Increases in discharge within the Red River watershed during the mid-1990s yielded higher loads of phosphorus to, and the subsequent eutrophication of Lake Winnipeg (McCullough et al. 2012). The elevated rates of primary production yield labile algal biomass that supplements the allochthonous OM respired in the water column during bloom senescence (Yezhova et al. 2021) but does not persist downstream to the lower Nelson River as a particulate OM source (Stainton 2019). Based on the results in this study, there does not appear to be metabolic coupling of the lower Nelson River and Lake Winnipeg. In contrast, the mean summertime NEP of 2007 in the Churchill River downstream of Southern Indian Lake was 0.067 ± 0.076 g C m^-2^ d^-1^ as estimated by D. Capelle (Freshwater Institute, Department of Fisheries and Oceans) using data from Stainton (2009). This is similar in magnitude to the unimpacted areas in this study, affirming the apparent lack of impact from impoundment at those locations and suggesting similar aquatic metabolism between the Nelson and Churchill rivers.

In a study by Teodoru et al. (2011) on the Eastmain-1 reservoir in boreal Quebec, prior wetlands showed some of the highest levels of benthic respiration and consequent CO_2_ emissions when compared to other pre-flood landscapes. However, the importance of pelagic respiration was not understated as it was a notable portion of total CO_2_ production within the reservoir (Teodoru et al. 2011). The results found here align with this paradigm as the backbay was the most heterotrophic location, but the disparity decreased once the respiration in the forebay was depth-integrated to account for the extent of pelagic processes. The La Romaine hydroelectric reservoir complex is a series of river impoundments in a cascade configuration with varying age, including La Romaine I (impounded 2016), La Romaine II (impounded 2014) and La Romaine III (impounded 2017). During 2017 and 2018, these reservoirs exhibited median NEP (GPP; R) values of -2.3 g C m^-2^ d^-1^ (0.86 g C m^-2^ d^-1^; -3.1 g C m^-2^ d^-1^), -1.5 g C m^-2^ d^-1^ (0.34 g C m^-2^ d^-1^; - 2.1 g C m^-2^ d^-1^) and -2.5 g C m^-2^ d^-1^ (0.43 g C m^-2^ d^-1^; -2.7 g C m^-2^ d^-1^), respectively (Barbosa et al. 2023). The mean rates of primary production in the KGS reservoir appear similar in magnitude to these median rates (< 1 g C m^-2^ d^-1^), despite the apparent lack of phosphorus limitation in the study area that is typically attributed to boreal water bodies (Schindler et al. 2008). In contrast, the median rates of respiration are greater and NEP is more heterotrophic in the Quebec reservoirs than mean values of the KGS, save for La Romaine II which is comparable to the backbay. The low degree of heterotrophy observed at unimpacted mainstem locations in the KGS study area aligns with the current understanding of carbon dynamics in lotic waters, wherein labile terrestrial OM decreases towards the downstream extent of the stream-river continuum resulting in lower mineralization rates (Hotchkiss et al. 2015; see Vonk et al. 2015).

### Limitations, caveats and future work

The combination of low productivity and dominant physical gas exchange processes yielded a low biological signal to noise ratio in [O_2_], and in one case (CL1) we were unable to interpret GPP and R values due to overwhelming presence of impossible estimates. Positive R estimates are generally considered a result of unconstrained physical processes (e.g., bubbles, Staehr et al. 2010; e.g., Hall et al. 2015) leading to error in k estimation, and the relatively low z_J at CL1 would have increased the sensitivity to error of the per unit time rate of gas exchange (k/z_J) used in calculation. Errors impacting GPP can result from non-linear responses to light (i.e., photoinhibition, light saturation) not accounted for in the linear relationship assumed, but developing a more complex model was beyond the scope of this study. There were multiple issues with the dataset for SL1, namely missing data due to instrument malfunction in 2020, estimation of the elevation profile due to lack of bathymetric data and the necessary correction to data in 2021. Statistical results were significantly sensitive to adjustment of SL1 [O_2_] input data, but not appreciably enough to change the main conclusions drawn here (Fig. S5). In light of these concerns, we have retained the data from CL1 and SL1 given the inconclusiveness of the temporal comparisons and downstream impacts, but have largely focused our analysis and discussion on results from within the KGS reservoir. Although data from Rabbit Creek and associated wetlands before impoundment are limited, it is established that inundation of terrestrial environments changes their carbon cycling dynamics (St. Louis et al. 2000). The resulting backbay itself is a new sub-environment resultant of reservoir creation and we have thus discussed it as “impacted” by impoundment here. However, we discourage interpreting the observed conditions as a fully attributable to impoundment effects without knowledge of baseline conditions. Future work could include the usage of bottle incubations or steady state isotope estimation (e.g., Barbosa et al. 2023) of metabolism to achieve a better understanding of spatial variability and the boundaries between sub-environments. Study of OM biodegradability or internal processes (e.g., photosynthetic efficiency) could confirm or elaborate some of the more speculative points made here.

## Conclusion

Based on the statistical analysis of the data gathered here and how it compared to prior research, we were able to broadly characterize aquatic metabolism in the study area with several preliminary conclusions. The inconclusive temporal comparisons and significant spatial differences reveal considerable variability in metabolic rates within the KGS reservoir, and that the impact of impoundment is not consistent throughout. The contrast between mainstem and backbay sub-environments appears to be determinant of aquatic metabolism, with pelagic and benthic respiration being of greater importance in each, respectively. Although both locations impacted by impoundment exhibited elevated primary production relative to other studied locations, respiration rate was greater in the backbay and it was the most heterotrophic sub-environment as a result. We propose HRT, allochthonous loading of OM and prior landscape qualities as key variables that may define this difference in aquatic metabolism. Light limitation appears to be the prevailing condition in the study area, which is consistent with previous research on the greater Nelson River system. As such, we propose that considering how environmental (e.g., climate) and anthropogenic influences on turbidity interact may provide insight into primary production within the KGS reservoir. Despite the unique water chemistry of the lower Nelson River and its source waters, the magnitude and patterns of aquatic metabolism were overall similar to those observed in other contemporary boreal reservoirs, if not lower in respiration rates. This assessment of aquatic metabolism in the lower Nelson River provides an initial understanding of major processes influencing GHG dynamics, and will inform the approach and interpretation of forthcoming research within a region of longstanding hydroelectric development.

## Acknowledgements

We thank Bailey Baldwin, Nicholas Decker and Yekaterina Yezhova for their assistance in receiving and processing water chemistry samples. Dr. Kristina Brown (UM) and Dr. Greg McCullough (UM) provided valuable feedback on the study design and interpretation. We also thank Wil DeWit (Manitoba Hydro) and other members of PEMP for collecting and providing the data used in this study, as well as for supporting our sampling program during the COVID-19 pandemic. The KGS is located on the traditional territory of the Tataskweyak Cree Nation, War Lake First Nation, Fox Lake Cree Nation and York Factory First Nation. It is these First Nations and Manitoba Hydro that form the Keeyask Hydropower Limited Partnership, and we thank them for permitting the use and publication of the data presented in this study.

## Author contributions

D.P.G. and T.N.P. conceived the study idea. D.P.G. performed data analysis and prepared the manuscript with draft review by T.N.P. A.D.S. coordinated water quality sampling efforts and acquisition of continuous monitoring data. R.G. liaised with collaborators at Manitoba Hydro for data sourcing. All authors reviewed the final manuscript.

## Funding information

This work was supported by the Natural Sciences and Engineering Research Council of Canada under Discovery Grant RGPIN-2019-06736 awarded to T.N.P.; and Manitoba Hydro funding awarded to T.N.P.

## Disclosure statement

R.G. is employed by, and funding for this study was provided by Manitoba Hydro, an entity that may benefit or be at a disadvantage due to the findings presented in this article or their implications.

## Data availability statement

The authors confirm that the original data generated in this study are available on the Canadian Watershed Information Network (CanWIN) at https://doi.org/10.34992/tsy9-7j16 and https://doi.org/10.34992/jgde-s608.

## Notes

https://doi.org/10.34992/tsy9-7j16

https://doi.org/10.34992/jgde-s608

