## Supplementary Material for "Aquatic metabolism throughout impoundment of a low productivity boreal reservoir using free water oxygen"

**Supplementary Information**

*Temporal extent of metabolism estimates*

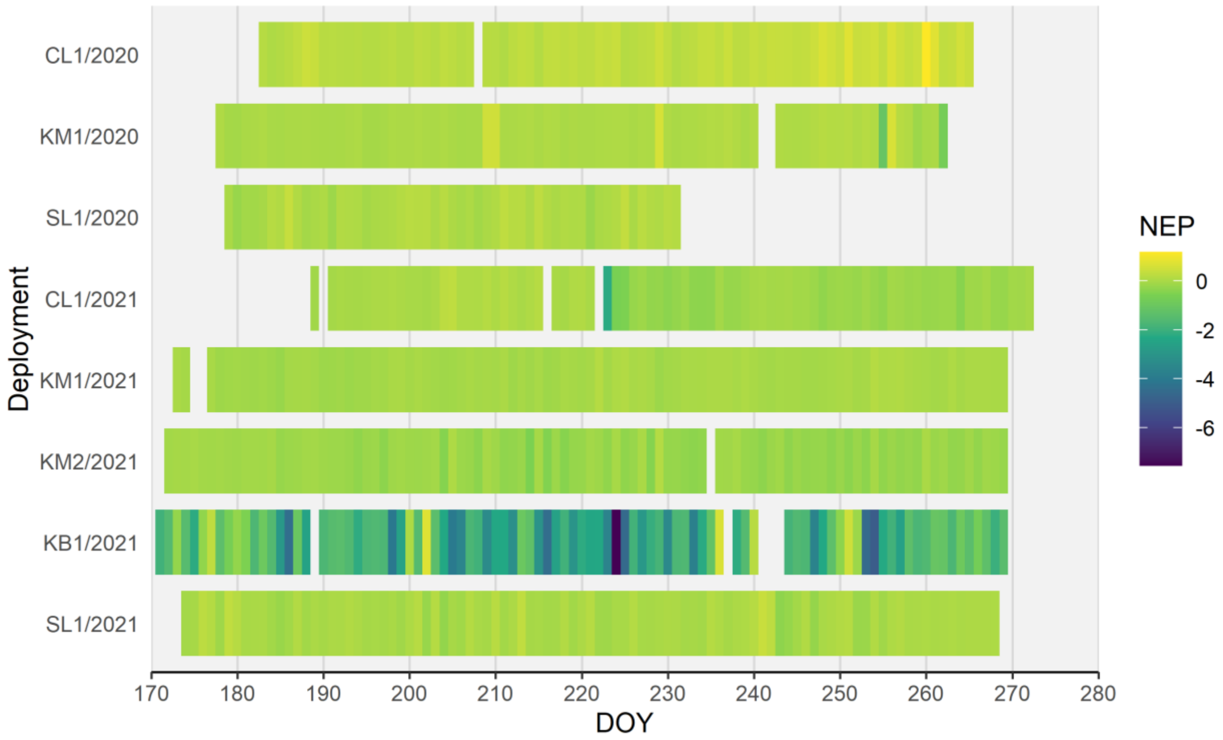


**Figure S1.** Availability of aquatic metabolism estimates by raft deployment (site/year) near and within the Keeyask Generating Station reservoir in northern Manitoba, Canada. The daily mean volumetric net ecosystem production (NEP in mg $O_2$ $L^{-1}$ $d^{-1}$) is depicted in gradient fill. Table 1 (main text) summarizes the metadata for each ID.

*Comparison to $CO_2$-based metabolism estimates*

The estimates of metabolism produced using free water oxygen curves were compared to estimates produced using the partial pressure of carbon dioxide ($pCO_2$) to corroborate the values. Continuous measurements were taken using a submersible $pCO_2$ sensor (CO2-Pro CV, Pro-Oceanus, Halifax, NS) on the floating raft deployed at KM1 (July 28 to September 9, 2020) and KB1 (June 26 to September 27, 2021). Measurement frequency was every four hours in 2020 and every eight hours in 2021. The model structure used was equivalent to the “bookkeeping” model described in Winslow et al. (2016) in terms of molar carbon quantities, which resulted in a daily mean value of net ecosystem production (NEP). To convert to mass-based units in terms of oxygen (mg $O_2$ $L^{-1}$ $d^{-1}$) as used by the *LakeMetabolizer* R package, photosynthetic quotient = 1.2 = respiratory quotient $^{-1}$ was used as in the main text and closely reflects Yezhova et al. (2021). The gas transfer velocity and $CO_2$ flux was calculated using a similar approach as $O_2$ described in the main text after Vachon and Prairie (2013) and scaled based on water temperature (Jähne et al. 1987; Raymond et al. 2012). Atmospheric partial pressure of $CO_2$ was assumed to be 405 µatm and solubility of $CO_2$ was calculated after Weiss (1974). The overall distribution of NEP estimates calculated via $O_2$ and $CO_2$ during the same time period were compared at each raft site using a Welch’s t-test and were found to be significantly different ($P < 0.001$; Figure S2). The lower mean NEP observed in the $CO_2$ approach versus the $O_2$ approach is similar to what Hanson et al. (2003) observed, who attributed the underestimation of gross primary production in lakes with pH > 8 to the influence of carbonate equilibria. Further, there are additional drivers of $pCO_2$ (e.g., carbonate weathering) that may introduce error into metabolism estimates based on $pCO_2$ (see Marcé et al. 2015). Linear regression was used to assess the comparability of each technique on a daily basis, which indicates that daily estimates of NEP were closer to 1:1 proportionality in the backbay than in the main channel (Figure S3). As such, it appears that errors occlude the biological [$O_2$] signal in the mainstem KM1 location more than in the backbay KB1 location. These results suggest the comparability of $O_2$-based metabolism estimates but indicate that care must be taken in interpreting or extrapolating results in mainstem sites.

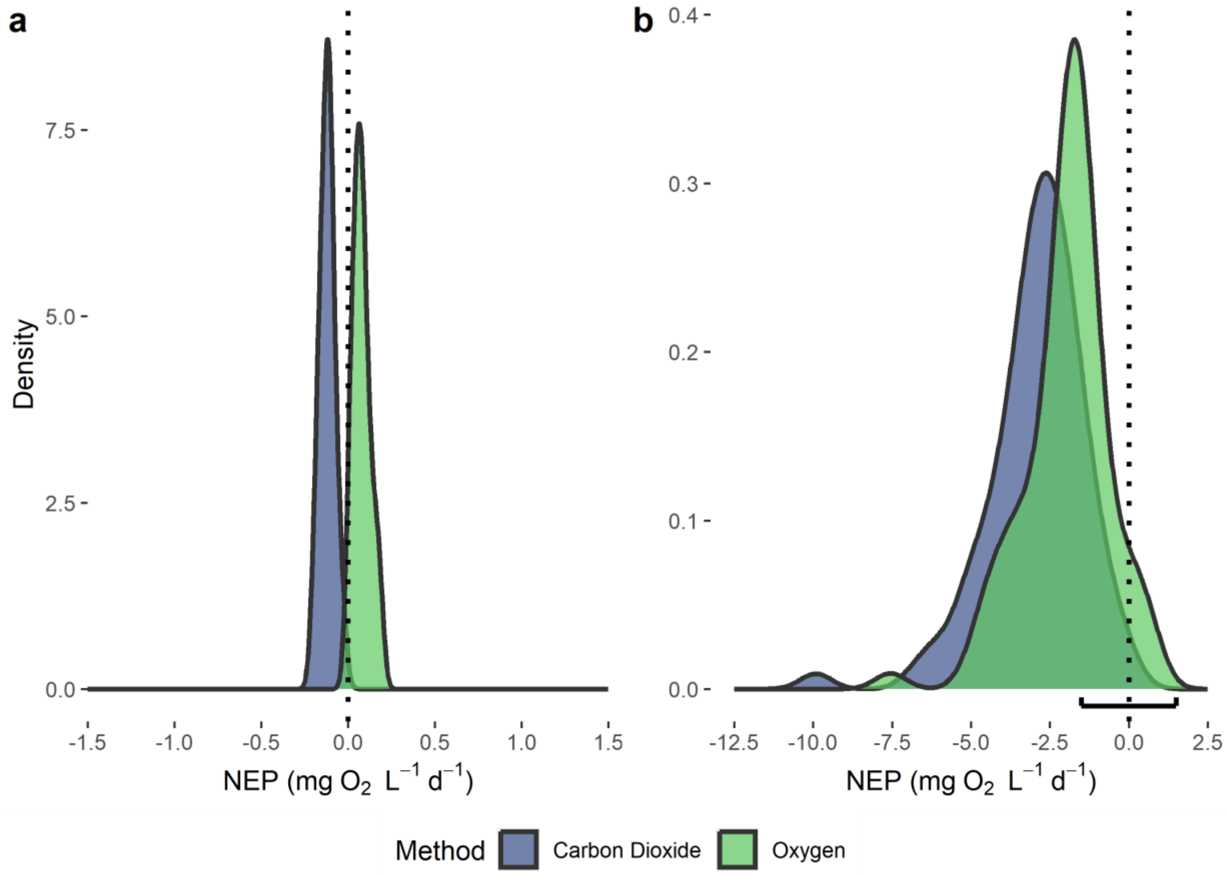


**Figure S2.** Density plots of net ecosystem production estimates calculated using diel $O_2$ and $CO_2$ curves at KM1 (a; 2020) and KB1 (b; 2021) sites of the Keeyask Generating Station reservoir in northern Manitoba, Canada. The black bracket on the x-axis in panel b indicates the range of the x-axis in panel a.

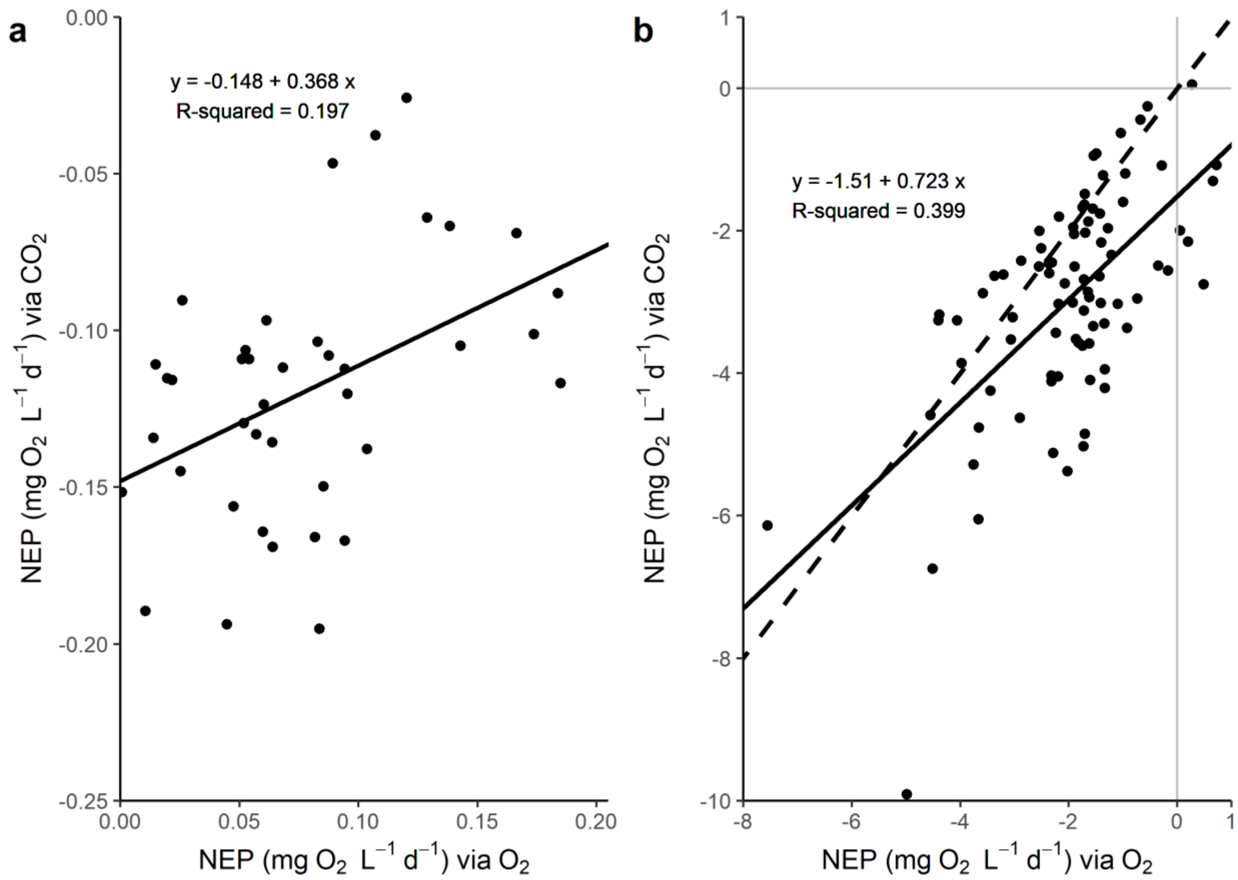


**Figure S3.** Scatterplots of net ecosystem production estimates calculated using diel [$O_2$] curves versus diel [$CO_2$] curves in the main channel (a; 2020) and backbay (b; 2021) sites of the Keeyask Generating Station reservoir in northern Manitoba, Canada. Linear regression line (solid) and equation are presented, as well as a 1:1 line (dashed) in plot b.

*Data correction for SL1 and sensitivity analysis*

Continuously measured $[O_2]$ at SL1 in 2021 was consistently high relative to discrete $[O_2]$ profiles (0.01 mg $L^{-1}$ resolution both), which indicated an error in calibration of the sensor (summarised in Table S1). To address this, the raw continuous $[O_2]$ data was conservatively adjusted by subtracting a nominal 1 mg $L^{-1}$ rather than e.g., the mean difference (1.339 mg $L^{-1}$) to avoid overcorrection given possible sampling or instrumental error in the discrete values. There were two concerns as a result of this data quality issue: (1) the sensitivity of SL1 statistical results to changes and the accuracy of conclusions as a result, and (2) the impact that including potentially inaccurate data from SL1 has on the results of other sites. To assess the first concern, two additional temporal Welch's t-test comparisons to 2020 data and two additional spatial Welch's ANOVA tests with Games-Howell post-hoc analysis were performed with SL1 metabolism estimates calculated using $[O_2]$ data reduced or increased by 0.5 mg $L^{-1}$ and compared to the original results in the main text. Adjustment resulted in either maintained significantly lower volumetric NEP (mg $O_2$ $L^{-1}$ $d^{-1}$) in 2021 (-0.5 mg $L^{-1}$, $P$<0.001) or a non-significant difference between 2020 and 2021 (+0.5 mg $L^{-1}$, $P$=0.55), which remains consistent with the conclusion that downstream impacts are uninterpretable or negligible (Figure S4). Although the results of Welch's ANOVA tests were unaffected ($P$<0.001), the spatial comparisons appeared more sensitive to adjustment of the $[O_2]$ data, with 0.5 mg $L^{-1}$ reduction in $[O_2]$ making areal NEP (g C $m^{-2}$ $d^{-1}$) comparable to KM2 (i.e., the forebay) versus a 0.5 mg $L^{-1}$ increase in $[O_2]$ making areal NEP making SL1 significantly more autotrophic than any site (Figure S5). This sensitivity is increased due to the relatively large mean depth at SL1 ($\bar{z}_{2021}$ = 18.1 m). Comparison of NEP at SL1 to other sites is invalid as a result, and thus additional data analyses (i.e., R and GPP comparison and coupling assessment) and discussion are centered on locations within the reservoir instead. To assess the second concern, spatial Welch's ANOVA tests with Games-Howell post-hoc analysis comparing volumetric and areal NEP were performed without data from SL1. The results of each Welch's ANOVA between sites remain significant when SL1 is excluded ($P$<0.001 both). The only resultant difference in spatial multiple comparisons by excluding SL1 is that the volumetric NEP of KM1 and CL1 become marginally significantly different ($P$=0.046) per the Games-Howell post-hoc test. As this does not conflict with the main conclusions drawn in this study, we have opted to retain SL1 for reference in statistical tests and figures.

**Table S1**. Comparison of discrete dissolved oxygen concentration profile data to continuously measured data at SL1.

| | | | *Discrete* | | | *Continuous* | | | |
|---|---|---|---|---|---|---|---|---|---|
| **Date** | **First Point** (CDT) | **Last Point** (CDT) | **Mean ± SD** (mg $L^{-1}$) | | | **Mean ± SD** (mg $L^{-1}$) | | | **Difference** (mg $L^{-1}$) |
| 7/6/21 | 09:17 | 09:30 | 9.264 | ± | 0.0089 | 10.616[a] | ± | 0.034 | 1.352 |
| 8/9/21 | 10:14 | 10:20 | 9.083 | ± | 0.0180 | 10.304[a] | ± | 0.475 | 1.221 |
| 9/26/21 | 13:56 | 13:58 | 9.987 | ± | 0.0076 | 11.429[b] | ± | 0.003 | 1.442 |

[a]Continuous mean value calculated using data from 20 min preceding first discrete point until 20 min after last discrete point

[b]Continuous mean value calculated using 60 min of data preceding sensor removal at 13:00 CDT

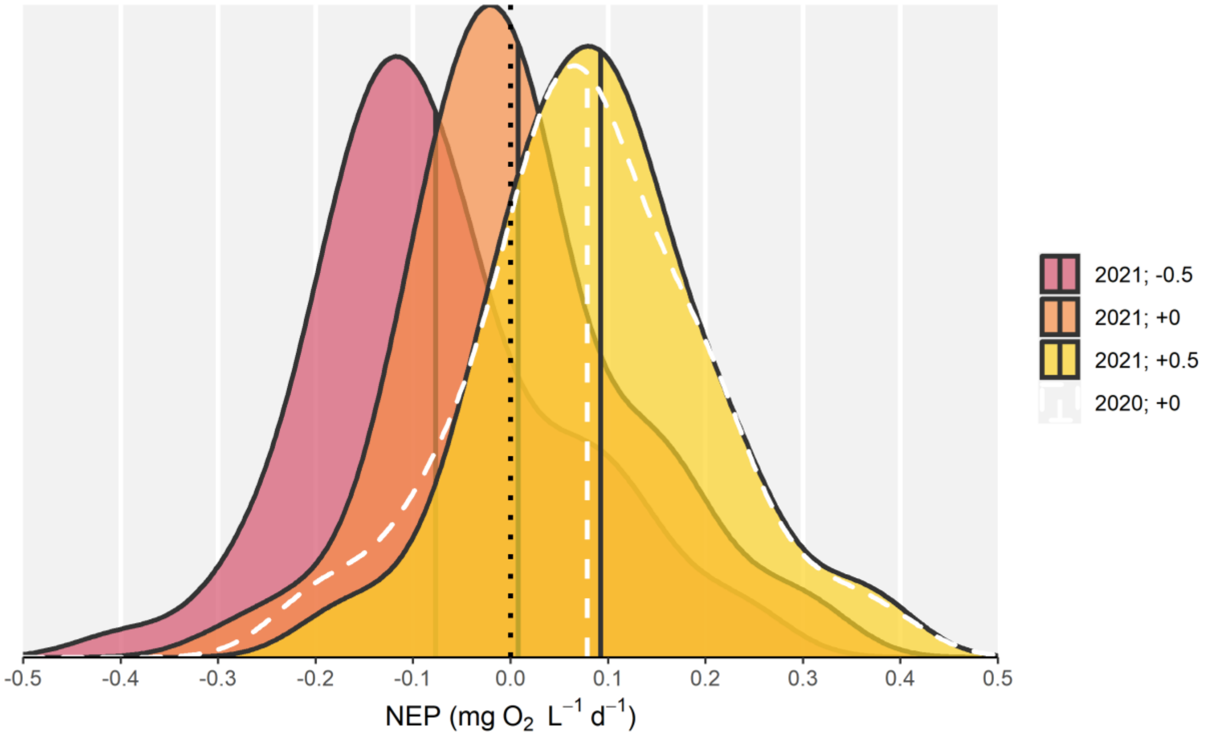


**Figure S4.** Density plots of volumetric net ecosystem production (NEP in mg $O_2$ $L^{-1}$ $d^{-1}$) at SL1 by adjustment to dissolved oxygen concentration data (mg $L^{-1}$) downstream of the Keeyask Generating Station reservoir in northern Manitoba, Canada. The mean of each distribution is represented by a vertical line. Data presented in this figure is from days before day-of-year 232.

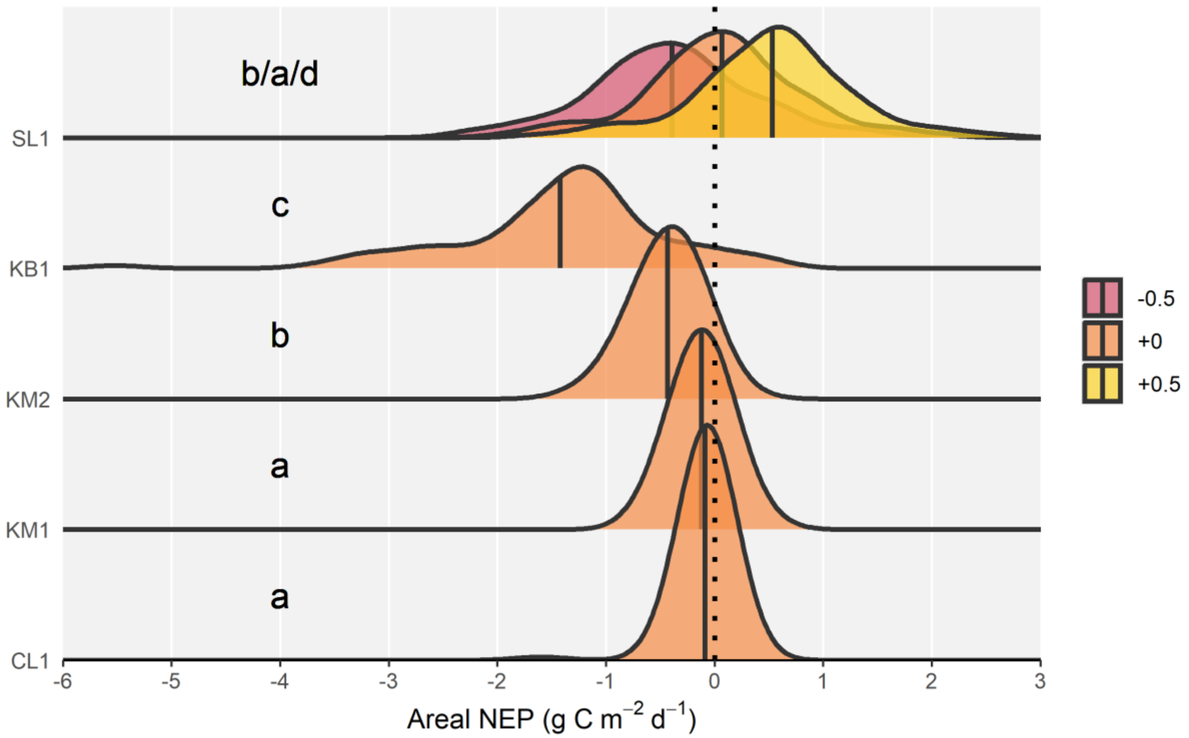


**Figure S5.** Density plots of areal net ecosystem production (NEP in g C $m^{-2}$ $d^{-1}$) by site and adjustment to dissolved oxygen concentration data (mg $L^{-1}$) near and within the Keeyask Generating Station reservoir in northern Manitoba, Canada during 2021. The mean of each distribution is represented by a vertical solid line. Results of Games-Howell post-hoc test are presented using compact letter display. Table 1 (main text) summarizes the metadata for each ID.

*References*

Hanson, P.C., Bade, D.L., Carpenter, S.R. and Kratz, T.K., 2003. Lake metabolism: Relationships with dissolved organic carbon and phosphorus. *Limnology and Oceanography*, *48*(3), 1112-1119. https://doi.org/10.4319/lo.2003.48.3.1112

Jähne, B., Heinz, G., and Dietrich, W., 1987. Measurement of the diffusion coefficients of sparingly soluble gases in water. *Journal of Geophysical Research: Oceans*, *92*(C10), 10767-10776. https://doi.org/10.1029/JC092iC10p10767

Marcé, R., Obrador, B., Morguí, J.A., Lluís Riera, J., López, P. and Armengol, J., 2015. Carbonate weathering as a driver of $CO_2$ supersaturation in lakes. *Nature Geoscience*, *8*(2), 107-111. https://doi.org/10.1038/ngeo2341

Raymond, P.A., Zappa, C.J., Butman, D., Bott, T.L., Potter, J., Mulholland, P., Laursen, A.E., McDowell, W.H., and Newbold, D., 2012. Scaling the gas transfer velocity and hydraulic geometry in streams and small rivers. *Limnology and Oceanography: Fluids and Environments*, *2*(1), 41-53. https://doi.org/10.1215/21573689-1597669

Vachon, D. and Prairie, Y.T., 2013. The ecosystem size and shape dependence of gas transfer velocity versus wind speed relationships in lakes. *Canadian Journal of Fisheries and Aquatic Sciences*, *70*(12), 1757-1764. https://doi.org/10.1139/cjfas-2013-0241

Weiss, R., 1974. Carbon dioxide in water and seawater: the solubility of a non-ideal gas. *Marine Chemistry*, *2*(3), 203-215. https://doi.org/10.1016/0304-4203(74)90015-2

Winslow, L.A., Zwart, J.A., Batt, R.D., Dugan, H.A., Woolway, R.I., Corman, J.R., Hanson, P.C. and Read, J.S., 2016. LakeMetabolizer: an R package for estimating lake metabolism from free-water oxygen using diverse statistical models. *Inland Waters*, *6*(4), 622-636. https://doi.org/10.1080/IW-6.4.883

Yezhova, Y., Capelle, D., Stainton, M. and Papakyriakou, T., 2021. Carbon fixation by the phytoplankton community across Lake Winnipeg. *Journal of Great Lakes Research*, *47*(3), 703-714. https://doi.org/10.1016/j.jglr.2021.03.003
